# Cheating (re)shapes pathogen virulence and antifungal resistance

**DOI:** 10.64898/2026.03.15.711879

**Authors:** H.M. Suraj, Frank P.J. Pieterse, Marnix A.J. van de Sande, Eric Bastiaans, Alfons J.M. Debets, André Fleißner, Jan A.L. van Kan

## Abstract

Filamentous fungi grow as fused, multinucleate networks that share secreted public goods vs private goods. We asked whether this sharing enables “cheater” nuclei to increase in frequency by exploiting goods produced by other nuclei, and whether such social conflict shapes virulence and antifungal resistance. We tested this in the gray mold pathogen *Botrytis cinerea* by contrasting an extracellular detoxification trait, a public good (enzymatic hydrolysis of the tomato saponin alpha-tomatine) with an intracellular antibiotic resistance trait, a private good (hygromycin phosphotransferase). In pairwise competitions, tomatinase-deficient nuclei gained advantage when rare against a constitutive producer, both *in vitro* and *in planta*, even though producers drive lesion expansion. An ordinary differential equation model fitted to the competition outcomes identified antibiotic gradients as the key driver of frequency-dependent selection and predicted stable coexistence of producer and non-producer nuclei across multinucleate bottlenecks. Cheating within fungal syncytia can therefore decouple virulence from reproduction and buffer the selection on antifungal resistance.

## Introduction

Cooperation is central to the evolution of multicellular organisms. It enables organisms to divide tasks, coordinate behaviour, and achieve outcomes that single units cannot reach alone. Many of these benefits rely on “public goods”, resources produced by the group for the collective benefit. But shared systems create vulnerability for exploitation. Individuals that use public goods without contributing can gain short-term advantages and weaken the performance of cooperative partners. These conflicts have been examined in several well-established systems especially in microbial communities and social amoeba (Smith & Schuster, 2019; Strassmann & Queller, 2011; Velicer & Vos, 2009), yet how they operate in multicellular eukaryotes such as filamentous fungi is still not well understood.

Filamentous fungi are not solitary microbes but complex social organisms that form multinucleate networks. These networks do not enforce strict cell boundaries, colonies routinely undergo germling and hyphal fusion (anastomosis), linking different parts of the mycelium (Fleißner et al., 2022). Fusion allows cytoplasm, organelles, and nuclei to freely move through the colony, creating a shared system where the nucleus (not the cell) acts as the cooperative unit (Roper et al., 2011, 2013). This connectivity brings benefits, including resource sharing and coordinated growth, but it also creates vulnerability for exploitation because non-cooperative nuclei can still gain access to shared resources and redirect them towards their own reproductive success. To limit this, fungi have evolved self/non-self-recognition systems, including allorecognition and heterokaryon incompatibility, which restrict fusion between genetically different individuals and help limit conflict within the mycelial network (Gonçalves et al., 2020). At the same time, traits that are private at the level of a single nucleus or hypha can effectively become public once cell fusion connects compartments and allows their products to spread through the colony. The evolutionary importance of these conflicts is illustrated in *Neurospora crassa*, where free fusion can favour the spread of cheater lineages that exploit somatic networks (Grum-Grzhimaylo et al., 2021). These fungal cheaters often exhibit fusion deficiencies. Despite their inability to initiate hyphal fusion, such cheaters passively parasitize fusion-proficient cooperators, benefiting from collective resources while contributing little to the shared somatic networks (Grum-Grzhimaylo et al., 2021).

In the plant pathogen *Botrytis cinerea*, infection depends on the secretion of “public goods” such as virulence factors and detoxifying enzymes that are shared within the colony. An example of such a public good is the secretion of the enzyme tomatinase, which is essential for pathogenicity on tomato plants (You et al., 2024). Tomato tissues accumulate α-tomatine, a potent steroidal glycoalkaloid which has a strong antifungal property, and thus provides basal resistance against pathogens (You & Van Kan, 2021). To overcome this chemical barrier, *B. cinerea* secretes a glycosyl hydrolase designated tomatinase, which detoxifies α-tomatine by converting it into the less toxic compound β_1_-tomatine (You et al., 2024). This enzymatic detoxification preserves fungal membrane integrity and is important for host colonization. Since tomatinase acts as a shared extracellular public resource, it creates the potential for exploitation by nuclear variants (“cheaters”) that contribute little to its costly production but still benefit from detoxified host tissue, thereby enhancing host colonization and reproductive success. To investigate these social dynamics, we utilized a wild-type strain of *B. cinerea* M3a isolated from grape (Quidde et al., 1999). Due to the insertion of a transposon inside the locus containing the genes required for tolerance to tomatine, M3a naturally lacks tomatinase (You et al., 2024). Thus, it is highly sensitive to α-tomatine and less virulent on tomato. We used wild-type M3a (WT) and a derived transformant (Bctom1; heterokaryon compatible) engineered to over-express both tomatinase (an extracellular enzyme) and hygromycin B phosphotransferase (a cytosolic enzyme). This experimental setup allowed us to contrast the extracellular tomatinase secretion as a cooperative “public good” (detoxifying α-tomatine for the colony as a whole) against hygromycin resistance as a “private good”, protecting only the cells expressing hygromycin B phosphotransferase that can detoxify the antibiotic hygromycin.

We performed *in-vitro* pairwise competition assays to determine whether the WT nuclei act as “cheater” (a nuclear variant not expressing tomatinase) in mixed cultures. Specifically, we investigated whether the wild type could benefit from the tomatinase provided by Bctom1 to gain a relative fitness advantage by not paying the full metabolic cost of enzyme production. Furthermore, we assessed the frequency-dependent nature of these interactions, aiming to distinguish the competitive dynamics of public versus private goods under varying starting frequencies. To test whether these *in-vitro* interactions translate to the complex environment in a host, we inoculated tomato leaves with single and mixed inoculum of wild-type vs Bctom1, and examined the competitive success *in-planta*.

To further explore the theoretical underpinnings of these dynamics, we employed ordinary differential equation modelling to develop a model that simulates cheating under varying selection pressures. Our model represents a simplified competitive environment with hypothetical strains ‘producers’ and ‘non-producers’, analogous to the Bctom1 and WT nuclei respectively. A sensitivity analysis with the model allowed us to identify which environmental and physiological parameters are key to the competitive success of cheaters. Additionally, we investigated how the multinucleate nature of *B. cinerea* spores might enhance the capacity of multiple nuclei to coexist in the population after generational bottlenecks. Through this combined experimental and theoretical approach, we aim to better understand how social conflicts constrain virulence and antifungal resistance, offering a potential explanation for the persistence of low-virulence strains in nature.

## Materials and methods

### Plant material

Tomato (*Solanum lycopersicum* cv. Moneymaker) was cultivated in a greenhouse (Unifarm, Wageningen UR) under 21/19 °C day/night conditions. The fungal inoculations were performed in the laboratory on detached tomato leaves.

### *B. cinerea* culture and plant inoculation

Culturing and spore production for the *B. cinerea* strain followed the protocols previously described by (You et al., 2024). Disease incidence was assessed on tomato leaves inoculated with 2,000 spores/spot in Gamborg’s B5 minimal medium (Duchefa), 15 mM sucrose, 10 mM potassium phosphate, pH 6.0). Leaves were incubated in high humidity under ambient light, and incidence was scored at 3 dpi.

### *In-vitro* and *In-planta* competition assay

Harvested spores of WT and mutant Bctom1 strains were adjusted to a concentration of 10^7^ spores/mL. Mixed spore suspensions were prepared at various ratios; for example, a 20:80 (WT : mutant) mixture was generated by combining 20 μL of WT and 80 μL of Bctom1 suspension with 900 μL of sterile distilled water. For *in-vitro* competition assay: a 2 μL droplet of the resulting mixture was inoculated at the center of Malt Extract Agar (MEA) plates. Plates were incubated at 20°C under a 16h light / 8h dark photoperiod. Spores were harvested and counted 7-dpi. For *in-planta* competition assay, the spore mixtures were inoculated on detached tomato leaves with 2,000 spores/spot in Gamborg’s B5 minimal medium, 15 mM sucrose, 10 mM potassium phosphate, pH 6.0, inside a transparent box with > 95% humidity. Following 24 h of incubation in high humidity/ambient light, leaves were transferred to a UV incubator (16:8 light:dark photoperiod) for 6 days, the spores were harvested and qPCR was performed to quantify the number of WT and mutant nuclei in the resulting spores. Competitive success of the cheater was calculated as the final frequency of the cheater after competition, divided by its initial frequency.

### qPCR for quantification of fungal DNA

DNA was isolated from spores using the Quick-DNA Fungal/Bacterial Miniprep Kit (Zymo Research; Cat. No. D6005). Total DNA concentration was determined using a NanoDrop spectrophotometer and adjusted to a final concentration of 5 ng/µL. For quantitative PCR (qPCR), 20 ng of template DNA was used per 25 µL reaction. Primers targeting the Tubulin A was used in-vitro experiments and Botcinic acid biosynthesis gene was used in in-planta experiments to quantify total fungal DNA, while primers specific to the hygromycin resistance cassette (hph) were used to quantify mutant DNA. qPCR was performed using the SensiFAST SYBR No-ROX Kit (Meridian Bioscience; BIO-98005) on a Bio-Rad CFX96 Real-Time System, following the manufacturer’s protocol. Primer sequences used for qRT-PCR in were: hygromycin resistance cassette (Hyg_Fw: TTAGCCAGACGAGCGGGTTCG; Hyg_Rv: GACGGTGTCGTCCATCACAGT), β-tubulin (*BcTubA*) (BcTubA_Fw: GGACGAGATGGAGTTCACTGAGG; BcTubA_Rv: TTGGGACCTCCTCTTCGTACTCC), and the botcinic acid biosynthesis gene *Bcboa6* (Bcboa6_Fw: GCAGGCTTGACTGGAGACTTG; Bcboa6_Rv: CAATGGTGTAGACGAGAACAGTC).

### Model description

To model the population dynamics within the mycelium, producer and non-producer nuclei were distinguished, each with their respective division equation dependent on the available resources (energy):

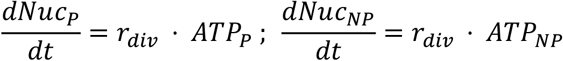

Where r_div_ is the nuclear division rate, and ATP_P_ and ATP_NP_ represent the (locally) available energy to the producer (P) and non-producer (NP) nuclei respectively. Both types of nuclei make use of a common pool of resources present in the intracellular space (E_int_), which is limited by the amount of external resources (E_ext_), which is defined at the start of simulation. The dynamics of the E_int_ is modelled with the following equation:

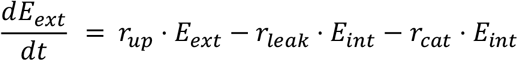

Where r_up_ represents the uptake rate, r_leak_ the leakage of resources out of the cell and r_cat_ the rate at which resources are catabolized in the mycelium. The dynamics of ATP_P_ and ATP_NP_ are then described by the equations:

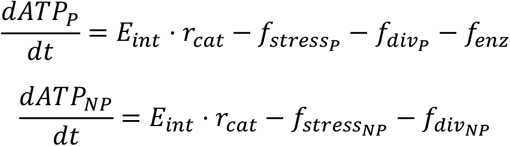

Where f_stress*_ represents the energy cost incurred by extracellular antifungal toxins, f_div_ represents the energy cost of division and growth, and f_enz_ represents the cost of producing the detoxifying enzymes. f_stress*_ is calculated differently for producer and non-producer nuclei:

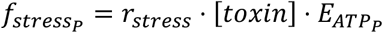

Where r_stress_ is a rate parameter describing the cost incurred relative to the concentration of the toxin ([toxin]). Critically, g_tox_ is a gradient, describing to what extent the non-producer nuclei experience more stress than the producers (due to diffusion processes limiting access to the detoxifying enzymes). The gradient is tied to the fraction of producer nuclei:

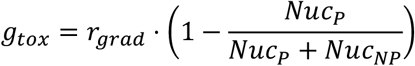

Where r_grad_ is a parameter describing the maximum ratio of stress experienced by non-producers. As the ratio of producer nuclei increases (and thus the amount of available detoxifying enzyme increases), g_tox_ decreases, reducing the additional stress cost experienced by non-producer nuclei.

### Parameter fit and sensitivity analysis

Model parameters were fitted to experimental data on the competitive success of several starting ratios. Parameters were manually adjusted until a good fit was found, and this parameter set can be found in Supplementary Table 1. With this fitted parameter set, we performed a sensitivity analysis where we varied one parameter at a time with ±25%. The mean absolute difference of the competitive ratio between a perturbed simulation and the baseline was used to rank the model parameters according to their sensitivity.

## Results

### Constitutive expression of tomatinase induces frequency-dependent selection

Previously, we generated a tomatinase-overexpressing strain (designated Bctom1) in *B. cinerea* M3a (WT) as described in You et al. (2024) and Figure 1A. The *niaD* locus was replaced with a cassette containing the hygromycin resistance cassette and an over-expressed tomatinase gene *Bctom*1 cloned from strain B05.10. We confirmed the functional impact of this modification by comparing the sensitivity of Bctom1 and the wild type to α-tomatine using spot assays. While both strains grew similar on control plates, the WT strain was severely inhibited by α-tomatine. In contrast, Bctom1 retained visible growth across all inoculum densities (Figure 1B), confirming that constitutive tomatinase expression confers tolerance to α-tomatine. This tolerance also resulted in increased virulence of Bctom1 compared to WT on tomato leaf (Figure 1D-E)

**Figure 1:**
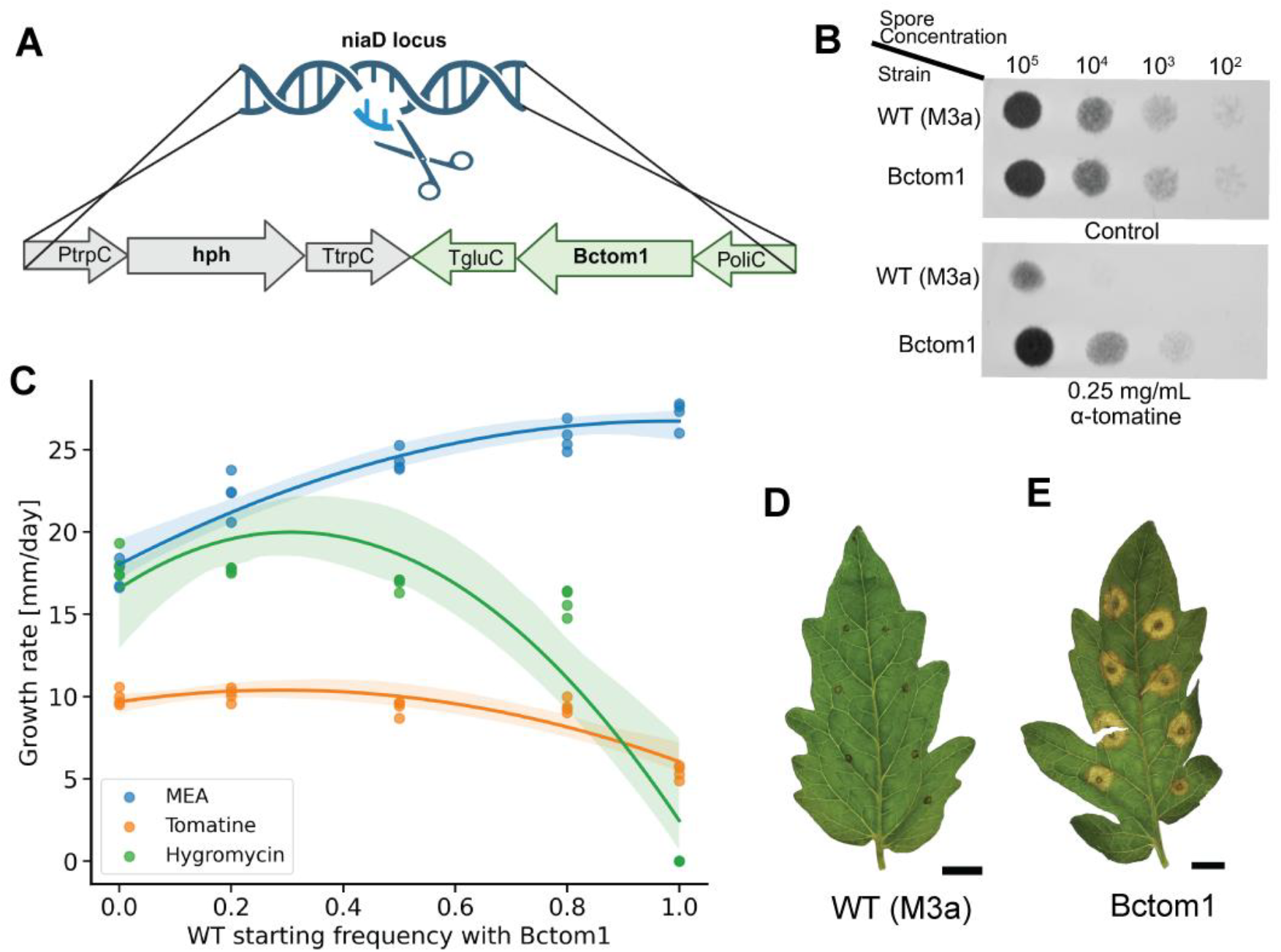
**A)** Scheme of the construct used to generate the tomatinase-overexpressing strain Bctom1. The cassette replaces the *niaD* locus in the M3a background and introduces the hph resistance marker together with the constitutively expressed tomatinas*e* gene. **B)** Sensitivity of wild-type and Bctom1 to α-tomatine. Serial dilutions of spores were spotted onto control BDES medium or medium containing 0.25 mg/mL α-tomatine. **C)** Radial growth rates of mixtures of wild type and Bctom1 across a range of starting frequencies on MEA, MEA supplemented with hygromycin (70µg/mL), or MEA containing α-tomatine (0.5 mg/mL). **D)** and **E)** WT and Bctom1 infected on the tomato leaf, photographed 3-dpi, scale bar is 1 cm.

We then assessed how the metabolic cost and social benefits of tomatinase secretion influence competition. The WT and Bctom1 strains were co-inoculated at various starting frequencies on three media: malt extract agar (MEA), MEA with hygromycin, and MEA with α-tomatine. The radial growth and spore yield were measured to capture overall fitness. On MEA, the mixed cultures displayed a unimodal growth pattern: growth rates (Figure 1C) and spore yields (Figure 2B) peaked when WT was abundant and declined as the frequency of the metabolic burden (in Bctom1) increased. Conversely, on media supplemented with hygromycin or α-tomatine, total colony spore yield dropped sharply as the frequency of the sensitive WT increased, reflecting the strong selection pressure against the non-resistant strain (Figure 2B).

**Figure 2:**
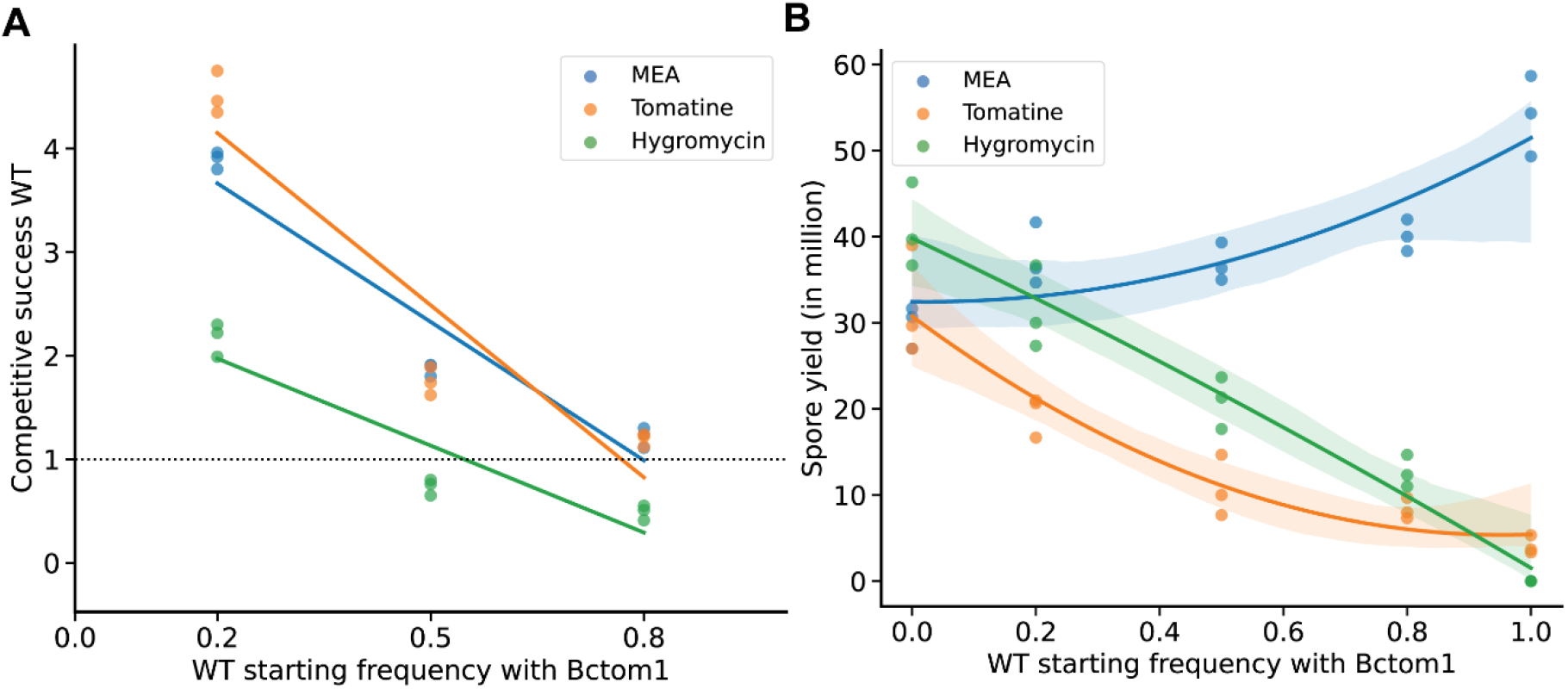
**A)** In vitro pair-wise competitions of wild type (WT) with Bctom1. The competitive success of WT was calculated using qPCR from the spores. **B)** Total spore yield (in millions) of WT and Bctom1 mixtures on MEA (blue), MEA supplemented with 70 ug/mL hygromycin (green) and MEA supplemented with 0.5 mg/mL α-tomatine. The areas around the trendlines are 95% confidence interval of the fitted curve. In both panels, the dots are the values from the three biological replicates.

To quantify competitive success, the relative fitness of each strain was calculated by comparing its final frequency to the initial frequency in the spore mixture used as inoculum. On MEA, the WT consistently outcompeted Bctom1 at all starting frequencies, revealing the significant fitness cost associated with constitutive tomatinase production (Figure 2A). In the presence of α-tomatine, this wild-type advantage was slightly higher in-comparison to MEA when WT was rare (starting frequency of 0.2). Under hygromycin selection, WT maintained a strong competitive advantage when rare, but the advantage disappeared at higher starting frequencies. Similarly, Bctom1 gained competitive advantage, only when it was rare under hygromycin selection (Figure 2A). This pattern demonstrates negative frequency-dependent selection in the presence of hygromycin, where WT benefits from the presence of Bctom1 at low frequencies but underperforms when it dominates in the population. Interestingly, these results show that non-producers, when rare, achieve much higher competitive success if the beneficial trait is a shared public good rather than when it is a private good (Figure 2A).

### WT acts as a cheater during mixed infection with Bctom1 on tomato leaves

In order to verify if the WT could act as cheater during *in-planta* infection similar to *in-vitro* inoculations, we performed competition assays between WT and Bctom1 with different starting frequency on tomato leaves. Disease incidence was positively correlated with the fraction of Bctom1 in the inoculum. Sole inoculation with Bctom1 resulted in high disease incidence (~85%), whereas increasing the frequency of WT significantly reduced disease severity (Figure 3A). In pure WT inoculations, leaves remained largely green and most primary inoculation spots non-expanding (Figure 3B). Despite the reduced disease severity in mixed infections (WT starting frequency of 0.8), WT nuclei displayed a reproductive advantage over Bctom1 within the lesions (Figure 3C). This suggests that while Bctom1 drives the necrotic infection, the WT nuclei confer the advantage of more efficient conversion of resources into spore production in a mixed population.

**Figure 3.**
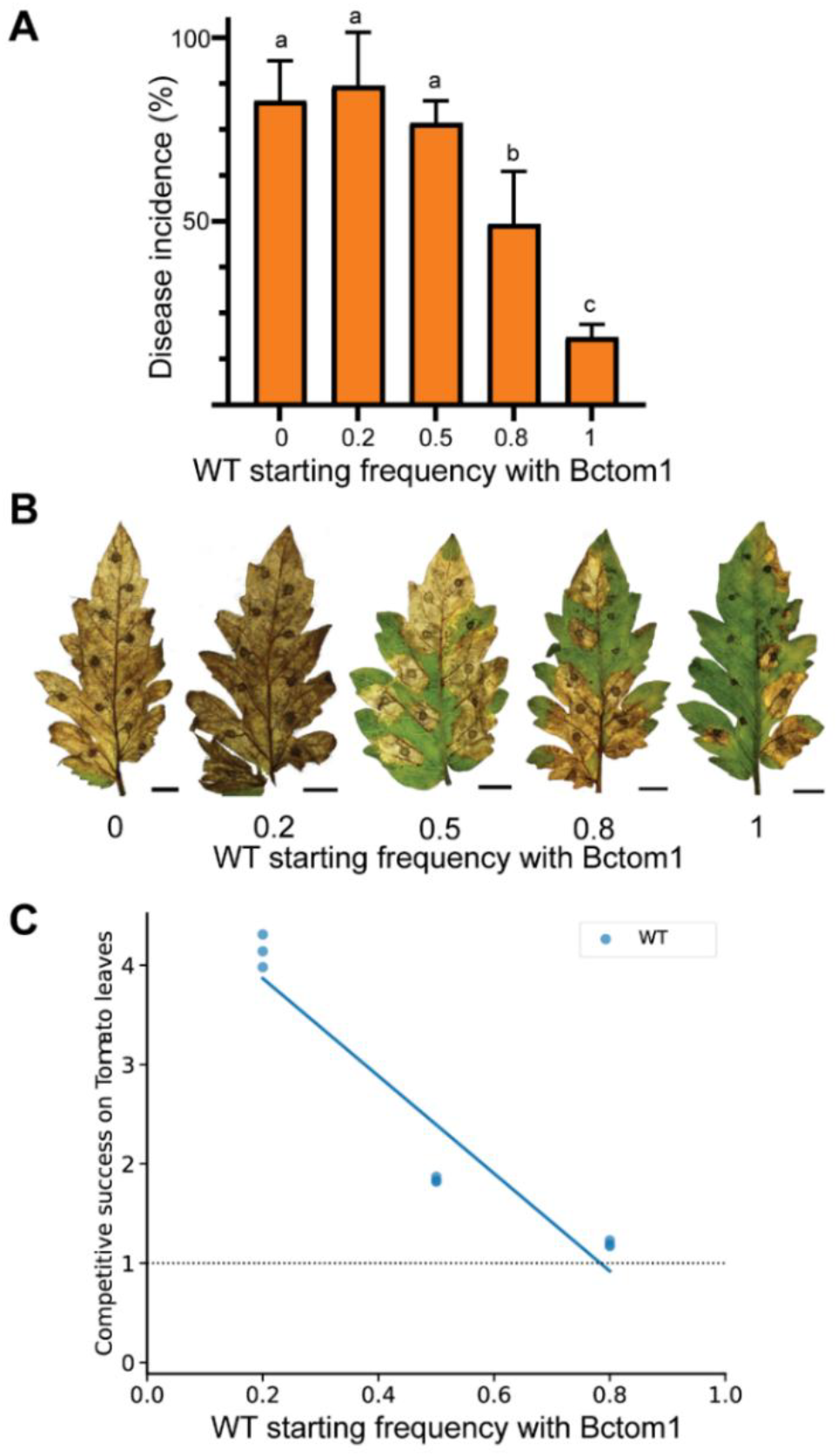
Frequency-dependent dynamics of WT versus Bctom1 competition on tomato leaves. **A**) Disease incidence measured at 3-dpi, quantified as the number of inoculation spots that develop expanding lesions. Incidence is high in populations dominated by Bctom1 but decreases significantly as the proportion of WT increases. Data represent mean incidence from 3 independent experiments; letters indicate significant differences (Chi-squared test, p-value < 0.01). **B)** Representative phenotypes of tomato leaves at 7 dpi. Necrotic severity correlates with the starting frequency of Bctom1. **C)** Competitive success of WT (blue) derived from spores harvested from infected tomato leaves.

### Antibiotic gradients drive cheating in a simplified model

To understand how non-producer (WT) nuclei gain a frequency-dependent advantage under antibiotic selection, we built an ordinary differential equation model with two nuclear types: “producers” (that contribute a protective detoxifying function at a cost) and “non-producers” (that do not pay this cost). As elaborated in the model description, the model simulates resource competition under constrained conditions. In the presence of an antibiotic, nuclei experience stress that can be relieved through enzymatic detoxification, but this detoxification carries a fitness cost for producer nuclei. We fitted the model to the measured competitive success of non-producers over a range of starting frequencies (Figure 4A). The fitted model reproduces the main pattern in the data: non-producers increase when rare (competitive success >1 at low starting frequencies) but lose the advantage as they become more common (competitive success <1 at higher starting frequencies). The non-producer nuclei benefit from the protection generated by producer nuclei when producer nuclei are common. To identify which model parameter controls this behaviour, we ran a sensitivity analysis by changing each fitted value by ±25%. The parameter representing the antibiotic gradient had the largest effect on predicted competitive success, shifting the curve most strongly, while similar changes in other terms produced smaller shifts (Figure 4B). The unequal antibiotic exposure is the main driver of cheating: steeper gradients strengthen the non-producer advantage when rare, while flatter gradients weaken it. Under a strong gradient, producer nuclei that face higher antibiotic loads pay the cost of protection and divide slowly, while non-producers in lower-stress regions divide faster yet still profit from shared protection. When stress is low (or the gradient is weak), this advantage shrinks and the producer cost dominates, so non-producers no longer gain. Two example experiments, with different starting ratios of producers and non-producers are visualised in Supplementary Figure 1. The time series of these simulations show how that cheaters have a competitive advantage as long as they are present in an environment where producers far outnumber the non-producers. However, when the non-producers strongly outnumber the producers, this competitive advantage flips around.

**Figure 4.**
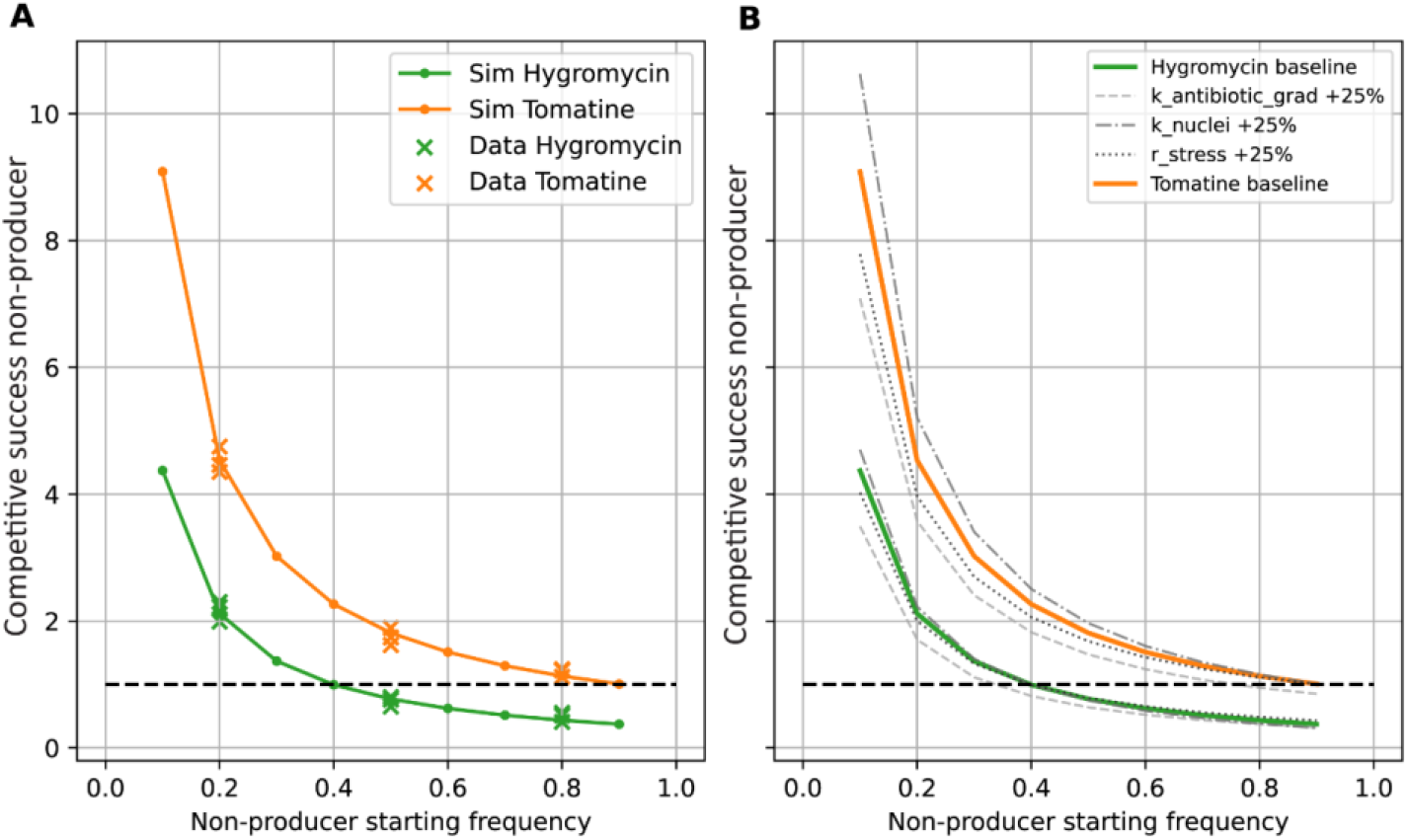
Experienced stress in combination with a local public detoxification enzyme enables cheating in modelling analysis. **(A)** Cheater model fitted to laboratory experiments of competitive success under different starting ratios. **(B)** Sensitivity of competitive success to the antibiotic gradient (k_antibiotic_grad), the ratio of ATP to nuclei (k_nuclei), and stress level (r_stress).

### Non-producer nuclei are stable in mixed cultures even as antibiotic pressure increases

Having defined a model that accurately describes outcomes of mixed cultures in a single cycle (from spore to sporulating mycelium), we asked how stable the fraction of producer nuclei is over multiple growth cycles. A scenario was investigated in which a bottleneck would allow only a few spores (and thus a few nuclei) to transfer to the next cycle. We randomly selected a number of nuclei from the end of the previous cycle to initiate a new cycle. After 30 cycles under continuous antibiotic pressure ([toxin]=1), the chance that the mycelium still contained two types of nuclei (heterokaryon) increased with the number of nuclei in the bottleneck (Supplementary Figure 2). This suggests that the multinucleate nature of fungal spores might allow multiple variant nuclei to exist within a mycelium across multiple growth cycles, which might provide benefits to the mycelium if overall fitness is higher in mixed populations.

We then asked how the concentration of antibiotic compounds in the environment might impact the stability of the heterokaryotic state. Both a linear increase (Figure 5A) and a linear decrease (Figure 5B) in antibiotic pressure were simulated, investigating the average fraction of producer nuclei at the end of each growth cycle, over 500 iterations. For these simulations we applied bottlenecks of 15 spores, as the heterokaryotic state of the mycelium was shown to be stable with this bottleneck size (Supplementary Figure 2). Paradoxically, the fraction of producers did not increase as antibiotic pressure increased 10-fold, suggesting that the positive selective pressure for producer nuclei is buffered by the frequency-dependent selection of non-producers. Remarkably, a decrease in antibiotic concentration did not lead to a decrease in producer nuclei, until antibiotic concentrations reached sufficiently low levels under which the production of detoxifying enzymes provided no further benefit. Thus, our model predicts that in mixed populations, non-resistant nuclei can stably coexist with resistant nuclei under a greatly varying range of antibiotic pressures.

**Figure 5.**
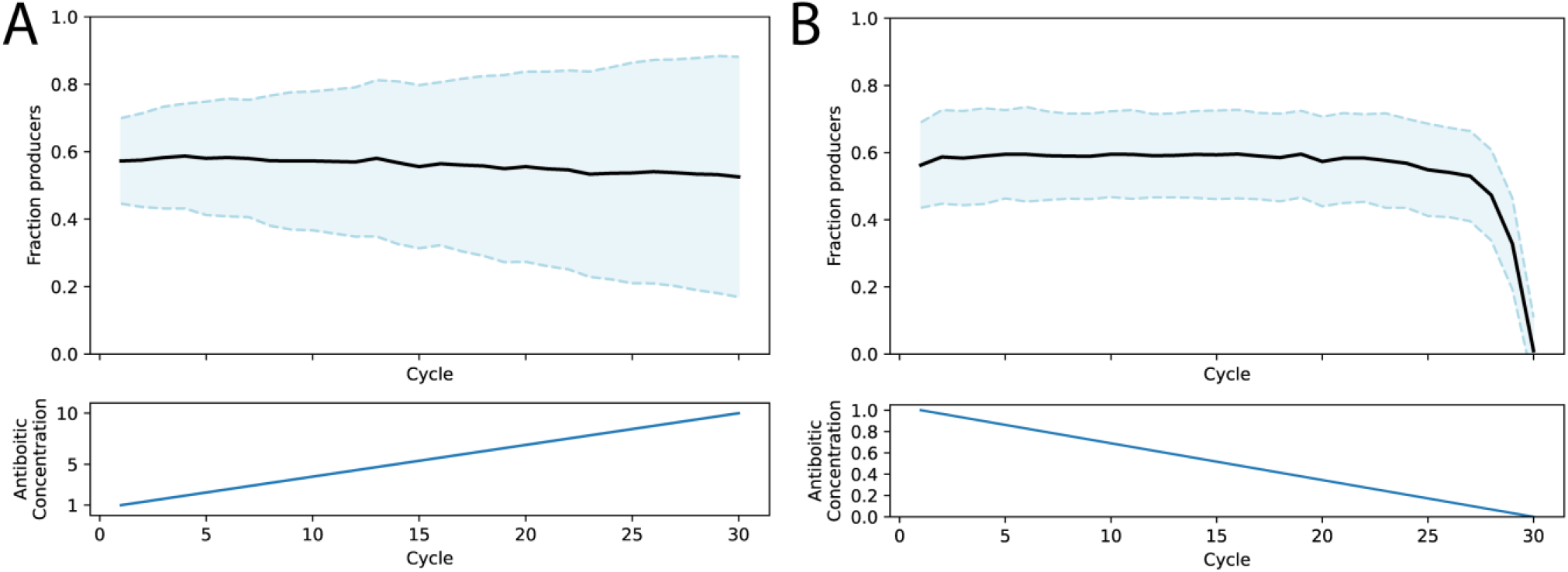
Stability of non-producers under varying antibiotic pressures. **A)** Average fraction of producer nuclei (top) over 30 cycles with increasing antibiotic concentration. Black line indicates average of 500 iterations with randomized generational bottlenecks of 15 nuclei. Blue area represents average ± standard deviation. **B)** Average fraction of producer nuclei (top) over 30 cycles with decreasing antibiotic concentration. Black line indicates average of 500 iterations with randomized generational bottlenecks of 15 nuclei. Blue area represents average ± standard deviation.

## Discussion

Filamentous fungi grow as connected, multinucleate networks where frequent hyphal fusion creates a shared cytoplasm. While this architecture supports collective traits vital for colony growth and host infection, it also creates vulnerabilities for exploitation. In *B. cinerea*, successful plant colonization requires the secretion of “public goods”. For example, the detoxifying enzyme tomatinase is secreted during infection of tomato plants to counteract the burden imposed by the host defence compound α-tomatine (You et al., 2024). This study demonstrates that nuclei lacking this enzyme can act as social cheaters, reaping the benefits of a detoxified environment without paying the metabolic costs of enzyme production. Crucially, a combination of *in planta* and *in vitro* experiments revealed that virulence and reproductive success can be decoupled in mixed infections. The tomatinase-producing nuclei (Bctom1) clearly drive the necrotic lesion expansion on tomato leaves, yet the non-producing nuclei efficiently parasitize this favourable environment, converting the shared resources into higher spore yields. This observation suggests that while disease symptoms track the presence of the public good producer, the reproductive success (spore yield) may disproportionately favour the non-producer. This dynamic is strikingly reminiscent of Defective Interfering Particles (DIPs) observed in viral infections (Huang & Baltimore, 1970; Vignuzzi & López, 2019). DIPs are naturally occurring viral mutants that lose essential genes (often for replication or structural proteins) but retain the ability to be packaged. They rely entirely on coinfection with a functional “helper” virus to propagate. Just as a high multiplicity of DIPs can lead to the collapse of a viral infection by draining resources from the helper, abundance of non-producing fungal nuclei reduces the collective output of mycelium, highlighting a universal vulnerability in shared biological systems.

It is important to note that we used a strong constitutive promoter in our experiments, whereas natural *B. cinerea* isolates carry the native *Bctom*1 promoter. In such isolates, tomatinase is only expressed in response to the exposure to α-tomatine (You et al., 2024), thus limiting the cost of production. Nevertheless, once a public good is secreted it can be exploited by non-producers. Previous reports have shown that tomato-infecting isolates of *B. cinerea* may carry a duplication or even triplication of the locus on Chromosome 8 containing the tomatinase gene *Bctom*1 and glycosyltransferase gene *Bcgt28a* (both contributing to tolerance to α-tomatine) (Mercier et al., 2021; Simon et al., 2022). Similarly, some *B. cinerea* isolates from grape lack these genes (Mercier et al., 2021) including strain M3a used in this study, highlighting the importance of such virulence factors in the interaction with specific host species. More broadly, virulence factors (including secreted enzymes and effectors) help pathogens colonize host tissues by weakening plant defense. In the future it would be interesting to test what happens during co-infection of two different pathogenic species on the same host, and whether public-good sharing creates frequency-dependent outcomes that shape pathogen population structure. Similarly, it would be interesting test two heterokaryon incompatible isolates of the same species to test if heterokaryon incompatibility reduces cheating on purely public vs private goods.

Previous studies in *N. crassa* showed that fusion-deficient mutants can act as cheaters by exploiting the somatic network of the wild type (Grum-Grzhimaylo et al., 2021). To investigate if this dynamic is conserved in *B. cinerea*, we assessed the social interaction between the WT and fusion-deficient mutant Δ*soft* (also known as Δ*Bcpro40*) (Haj Hammadeh et al., 2022). The Δ*soft* strain carries a hygromycin-resistance cassette at the *soft* locus, whereas WT is hygromycin-sensitive, which allowed us to test their interaction under selection. In contrast to *N. crassa*, the *B. cinerea* Δ*soft* mutant did not behave as a cheater. On non-selective MEA, WT outcompeted Δ*soft* (Supplementary Figure 4A). Under hygromycin selection, pure WT cultures failed to produce spores, but mixed cultures restored sporulation and produced more spores than the Δ*soft* monoculture (Supplementary Figure 4B). These results argue against one-sided exploitation and instead support reciprocal complementation, where the WT gains access to hygromycin resistance, whereas Δ*soft* gains from association with fusion-competent WT. Thus, goods that appear private at the level of an individual hypha can become shared once hyphal fusion links the colony. A more direct test of this hypothesis would be to compare heterokaryon-compatible and heterokaryon-incompatible producer and non-producer pairs, and investigate whether fusion is required for sharing of private goods but not for public goods. However, vegetative incompatibility in *Botrytis cinerea* remains comparatively underexplored at the molecular level. Despite the fact that the *B. cinerea* genome encodes several dozens of proteins that contain *het* domains, there is no functional evidence that any of these genes participates in heterokaryon (in)compatibility. Arshed et al. (2023) identified and functionally characteriezed the first vegetative incompatibility (*vic*) locus in *B. cinerea*, but their study highlighted experimental limitations, including the need for genetic mapping in sexual offspring and assay-dependent interpretation of incompatibility.

Unlike the private good scenario described above, tomatinase acts as a public good and therefore does not require direct fusion between producer and non-producer nuclei to be exploited. Consistent with this, competition assays performed with mycelium also showed that the non-producer (WT) already had a competitive advantage during vegetative growth against the producer strain (BcTom1) (Supplementary Figure 4). This suggests that, in *B. cinerea*, cheating on tomatinase begins during mycelial expansion, likely because non-producers avoid the cost of enzyme production while still benefiting from detoxification of the surrounding environment. To better understand this behaviour, we developed an ordinary differential equation model to examine how cost-free access to detoxified host tissue can promote the spread of cheaters under antibiotic pressure. The model highlights that unequal antibiotic distribution driven by spatial diffusion gradients is the primary driver of cheating. Non-producers that occupy lower-stress microenvironments created by producers can divide rapidly because they avoid the metabolic cost of enzyme synthesis. This highlights that environmental structure and population structure may therefore be a key determinant of whether, and how strongly, cheating occurs (Cremer et al., 2019; Testa et al., 2019; Velicer et al., 1998). For example, infection by the necrotrophic pathogen *B. cinerea* on tomato leaves triggers host cell death, releasing α-tomatine from the vacuoles and generating localized zones with high tomatine concentrations, effectively creating a structured environment during infection. It would be informative to test how different degrees of environmental structure shape population dynamics, using experimental setups that more closely resemble agricultural conditions.

The multi-generational modelling simulations revealed that the multinucleate nature of fungal spores could cause two types of nuclei to stably coexist across generational bottlenecks. When introducing this bottleneck in our model, we randomly sampled nuclei from the mycelium. We thereby assumed that the multinucleate spores of *B. cinerea* result from an influx of nuclei from the mycelium into the conidiophore, rather than through the division of a single founder nucleus. For *Aspergillus oryzae* it was shown that conidiospores form by active trafficking of nuclei rather than through division (Ishi et al., 2005). Glomeromycota provide an extreme example of multinucleate spores which can contain hundreds of nuclei. In *Glomus etunicatum* it was shown that such multinucleate spores form by recruitment of nuclei from the mycelium, which resulted in a lower mutational load in the population (Jany & Pawlowska, 2010). It would be interesting to study the mechanisms of sporogenesis in *B. cinerea*, and whether this has implications for its population dynamics and evolution. We observed that producers and non-producers can stably coexist across a wide range of fluctuating antifungal pressures.

From an agricultural perspective, this is highly consequential. It suggests that applying fungicides or relying on plant chemical defenses may not purge susceptible (non-producing) strains from the field. In cases where compatible isolates can fuse, nuclei from sensitive isolates may persist within shared hyphal networks by benefiting from resistance traits provided by other nuclei. More generally, for extracellular public goods, sensitive strains may also persist in mixed infections without requiring fusion. Such interactions may help explain why sensitive pathogen populations can rapidly rebound once chemical pressure is removed. Moreover, the presence of low-virulence nuclei in a population may not always decrease the virulence of heterokaryotic populations. We showed that when ≤50% of nuclei were non-producers (of the virulence-related enzyme tomatinase), the disease incidence was not affected. Interestingly, it has even been reported that the presence of low-virulence nuclei can sometimes even increase virulence at the population level (Lindsay et al., 2016). Testing mixed infections with compatible and incompatible strains would help determine how such interactions shape collective virulence. These results emphasize that fungal pathogens should be considered as interacting multinucleate populations rather than as isolated individuals.

**Supplementary table 1:** Optimised parameter set and initial conditions used for the model simulations.

| Name | Type | Value | Description |
| --- | --- | --- | --- |
| r_up | parameter | 0.09 | Uptake of glucose |
| r_out | parameter | 0.01 | Leakage or export of glucose |
| r_cat | parameter | 0.01 | Production of ATP from glucose |
| r_stress_hygromycin | parameter | 0.28 | Stress rate for hygromycin condition |
| r_stress_tomatine | parameter | 0.69 | Stress rate for tomatine condition |
| r_enzyme_prod | parameter | 0.01 | Rate of enzyme production |
| r_nuc_div | parameter | 0.001 | Rate of nuclear division in terms of ATP loss |
| r_detox | parameter | 0.0001 | Rate of detoxification by the produced enzyme |
| k_nuclei | parameter | 4.12 | The ratio of ATP to nuclei |
| k_antibiotic_grad_hygromycin | parameter | 2.58 | Max stress gradient in hygromycin condition |
| k_antibiotic_grad_tomatine | parameter | 1.12 | Max stress gradient in tomatine condition |
| k_nuc_to_vol | parameter | 1e-12 | Ratio betw N_nuc and the vol of one fungal cell, relative to total petri vol |
| gluc_ext | init condition | 20.0 | External glucose at start of simulation |
| gluc_int | init condition | 2.0 | Internal glucose at start of simulation |
| detox_enzyme | init condition | 0.0 | Available detox enzyme at start of simulation |
| antibiotic | init condition | 1.0 | Available antibiotic (stressor) at start of simulation |
| E_stored | init condition | 5.0 | Stored energy at start of simulation |
| nuclei | init condition | 10.0 | Total number of nuclei at start of simulation |
| frac_producers | init condition | 0.5 | Division of genotypes (producer vs.non-producer) amongst available nuclei |
| E_atp_p | init condition | 1.0 | ATP of producers at start of simulation |
| E_atp_np | init condition | 1.0 | ATP of non-producers at start of simulation |

## Supplementary figures

**Supplementary figure 1:**
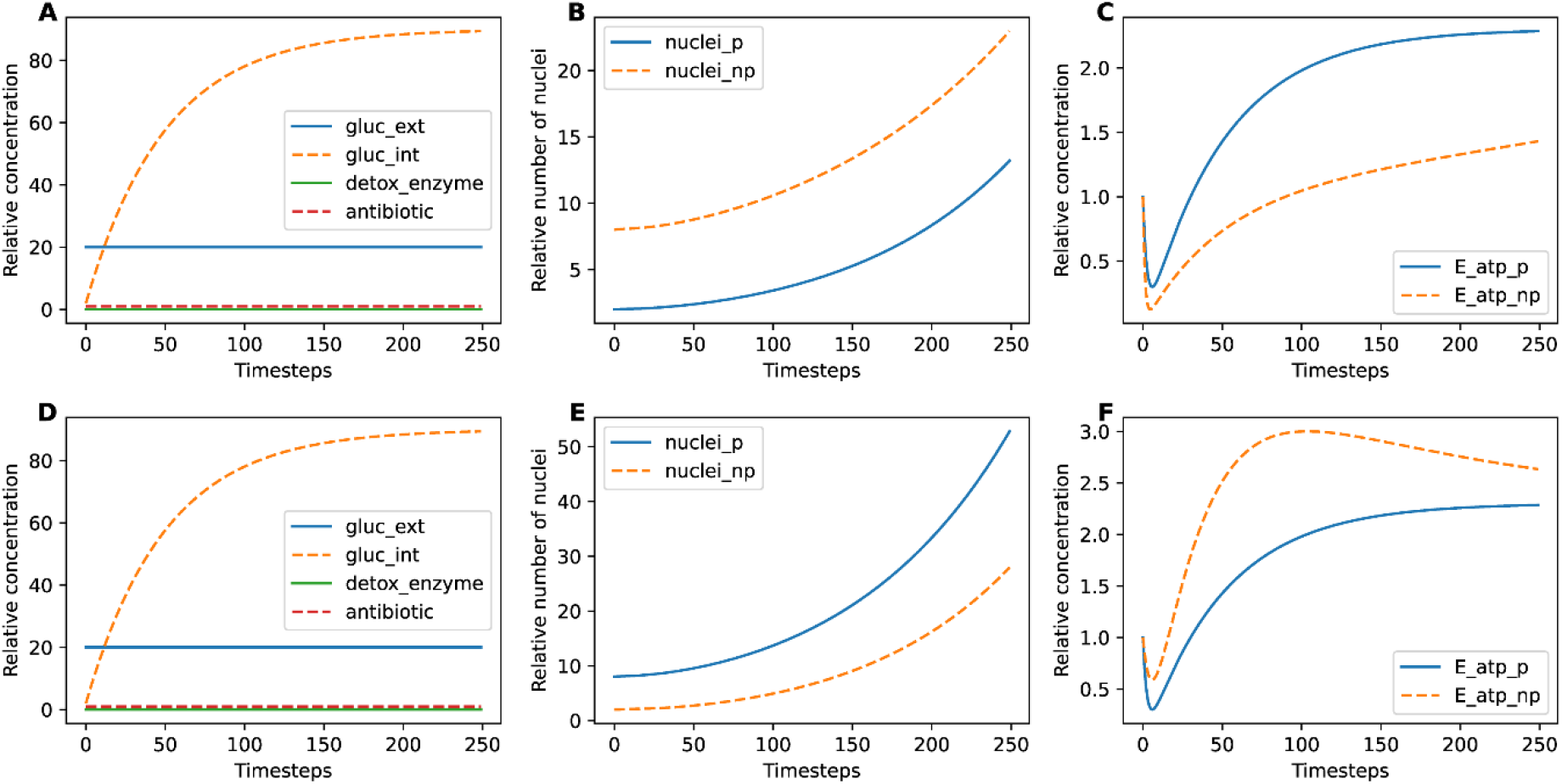
Time series of individual model simulations. Panels (A), (B) and (C) show a simulation where producers (analogous to Bctom1) have a starting ratio of 0.2 and the non-producers (analogous to WT) have a starting ratio of 0.8. Panels (D), (E), and (F) show a simulation where producers (Bctom1) have a starting ratio of 0.8 and the non-producers (WT) have a starting ratio of 0.2. The competitive success shows in for instance figure 5 is derived from the relative increase in nuclei (end compared to start of simulation) of the non-producers and producers, respectively.

**Supplementary figure 2:**
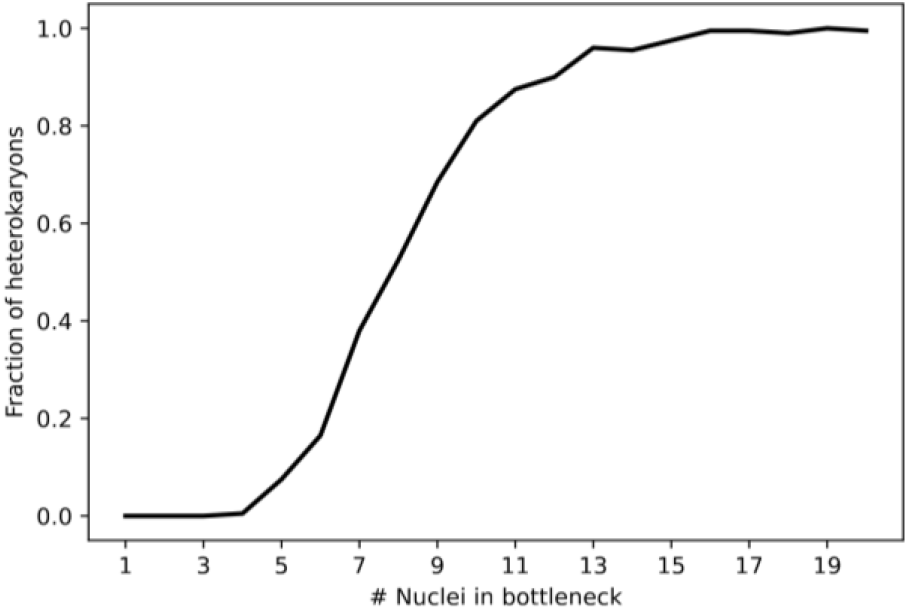
Fraction of heterokaryotic mycelia increases as more nuclei are selected in bottlenecks. Average number of heterokaryotic mycelia (containing both types of nuclei) is shown after 30 growth cycles under continuous antibiotic pressure ([toxin] = 1.0).

**Supplementary figure 3:**
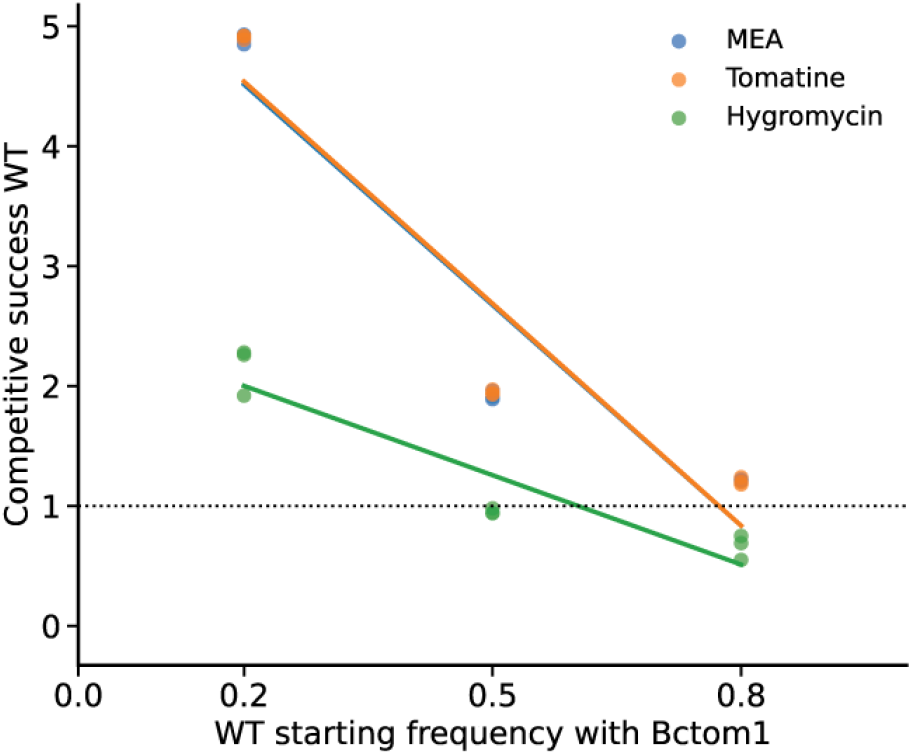
In pair-wise competitions of wild type (WT) with Bctom1 mycelium 3 days post inoculation. The competitive success of WT was calculated using q-PCR for the dna extracted from the mycelium.

**Supplementary figure 4:**
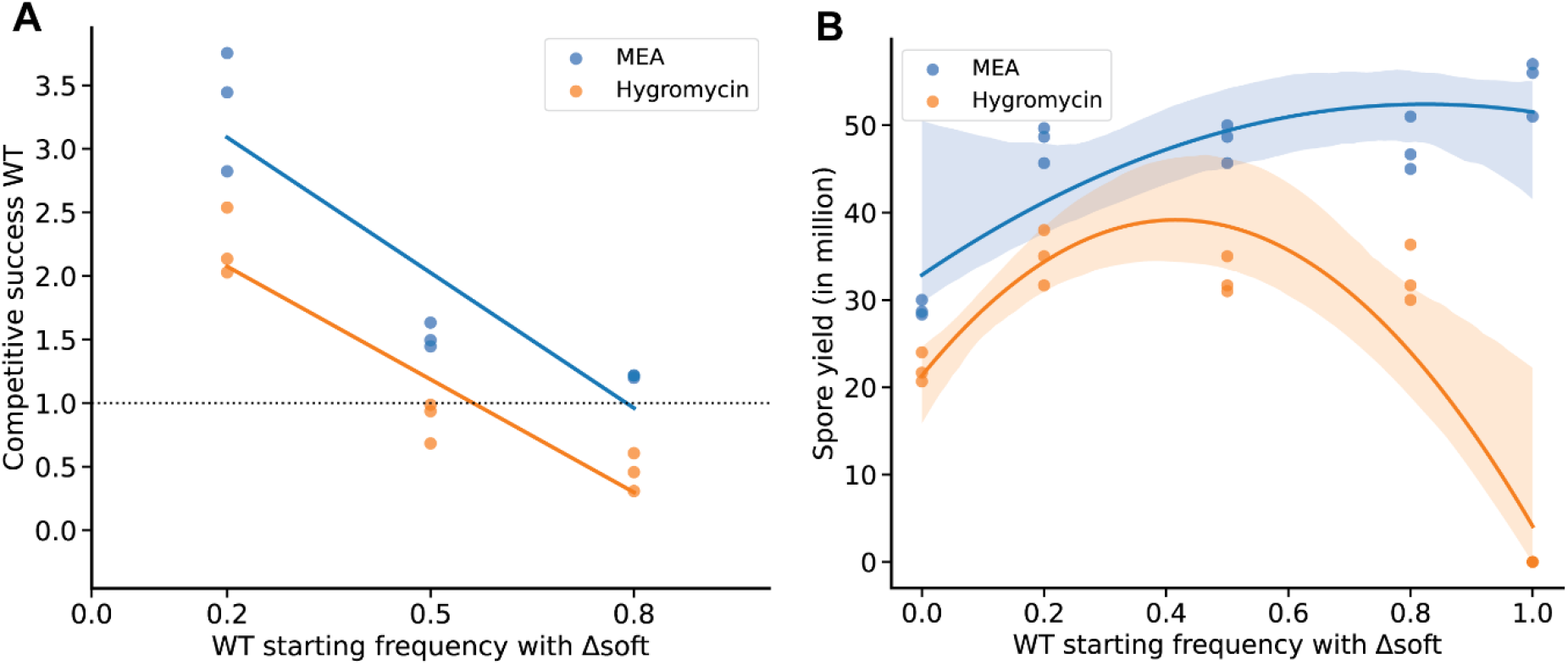
**A)** Competitive success of WT relative to Δ*soft* on MEA and MEA containing hygromycin (70 ug/mL). **B)** Total spore yield (in millions) of WT and Δ*soft* mixtures on MEA (blue) and MEA supplemented with hygromycin (orange). Shaded regions represent the 95% confidence interval of the fitted curve.

## Notes

### Competing Interest Statement

The authors have declared no competing interest.

### Funding

- TKI Green Genetics: LWV21.291 / TU-2021-20

